# Abscisic Acid modulates neighbor proximity-induced leaf hyponasty in Arabidopsis

**DOI:** 10.1101/2022.02.18.481016

**Authors:** Olivier Michaud, Johanna Krahmer, Florian Galbier, Maud Lagier, Vinicius Costa Galvão, Yetkin Çaka Ince, Martine Trevisan, Jana Knerova, Patrick Dickinson, Julian M. Hibberd, Samuel C. Zeeman, Christian Fankhauser

## Abstract

Leaves of shade avoiding plants like *Arabidopsis thaliana* change their growth pattern and position in response to a low red to far-red ratio (LRFR) that is encountered in dense plant communities. In LRFR, transcription factors of the phytochrome interacting family (PIFs) are de-repressed. PIFs induce auxin production, which is required to promote leaf hyponasty thereby favoring access to unfiltered sunlight. Abscisic acid (ABA) has also been implicated in the control of leaf hyponasty. In addition, gene expression patterns suggest that LRFR regulates the ABA response. Here we show that, in leaves, LRFR leads to a rapid increase in ABA levels. Changes in ABA levels depend on PIFs, which regulate the expression of genes encoding isoforms of the enzyme catalyzing a rate-limiting step in ABA biosynthesis. Interestingly, ABA biosynthesis and signaling mutants have more erect leaves in white light but respond less to LRFR. Consistent with this, ABA application decreases the leaf angle in white light but this response is inhibited in LRFR. Tissue-specific interference with ABA signaling indicates that an ABA response is required in different cell types for LRFR-induced hyponasty. Collectively, our data indicate LRFR triggers rapid PIF-mediated ABA production. ABA plays a different role in controlling hyponasty in white light compared to LRFR. Moreover, ABA exerts its activity in multiple cell types to control leaf position.

**One sentence summary:** Abscisic acid biosynthesis and signaling are required to promote leaf hyponasty in response to a drop in the red to far-red ratio.

## Introduction

Leaf movements are controlled by the interplay of circadian and light signals (McClung 2013; Hopkins et al. 2008; Dornbusch et al. 2014). The physiological significance of these movements is not fully understood, however, leaf position influences both photosynthesis and water loss (Hopkins et al. 2008). In open environments, light reaches intensities higher than the photosynthesis saturation point. Having more erect leaves diminishes light interception and favors cooling which is proposed to be beneficial (Hopkins et al. 2008). Consistent with this idea, Arabidopsis accessions from lower latitudes have more erect leaf angles (Hopkins et al. 2008). In addition to these diel leaf movements, several environmental cues including flooding, high temperature and shade lead to rapid upwards leaf repositioning (hyponasty) (Voesenek et al. 2006; Casal and Balasubramanian 2019). This movement is thought to help leaves acclimate to such unfavorable conditions for example by enhancing cooling capacity in warm environments (Crawford et al. 2012; Voesenek et al. 2006; Casal and Balasubramanian 2019).

At the molecular level the mechanisms underlying hyponasty are probably best understood in response to a decrease of the red to far-red ratio (low red/far-red, abbreviated LRFR here). LRFR occurs in dense vegetational communities and is a modification of the light environment experienced by plants prior to actual shading (Casal 2013; Ballare and Pierik 2017). Leaf hyponasty is induced by different elements of canopy shade including both low light and LRFR (Mullen et al. 2006; Millenaar et al. 2009; Moreno et al. 2009; Casal 2013; Ballare and Pierik 2017). The primary photoreceptor controlling LRFR-regulated responses is phytochrome B (phyB) (Casal 2013; Ballare and Pierik 2017). LRFR inactivates phyB leading to the de-repression of several Phytochrome Interacting Factors, particularly, PIF4, PIF5 and PIF7 (Casal 2013; Ballare and Pierik 2017; Cheng et al. 2021). These PIFs promote shade-induced elongation of the hypocotyl, leaf petioles, and leaf hyponasty (Keller et al. 2011; Kohnen et al. 2016; de Wit et al. 2016; Michaud et al. 2017; Pantazopoulou et al. 2017). phyB inactivation at the leaf rim leads to PIF-mediated auxin production in the blade, followed by auxin transport and redistribution in the petiole where it promotes upwards leaf repositioning (Michaud et al. 2017; Pantazopoulou et al. 2017; Gao et al. 2020).

Beside auxin, other hormones also contribute to environmentally-controlled leaf repositioning in Arabidopsis. Ethylene is particularly important to promote upwards leaf repositioning in response to waterlogging (Millenaar et al. 2009; Rauf et al. 2013). In contrast, ethylene is not required for low light-induced hyponasty and it negatively regulates high temperature-induced hyponasty (Millenaar et al. 2009; van Zanten et al. 2009). The role of abscisic acid (ABA) in controlling leaf position is also complex and poorly understood. ABA inhibits ethylene-induced hyponasty and in white-light grown plants genetic and pharmacological experiments indicate that ABA promotes downwards leaf repositioning (Mullen et al. 2006; Benschop et al. 2007). In contrast, ABA can also positively regulates hyponasty when plants are treated with an elevated temperature (van Zanten et al. 2009). Access to water is essential for growth and reversible cellular expansion, suggesting that ABA might be an important regulator of leaf positioning (Tardieu et al. 2015; Yoshida et al. 2019). Indeed, differential petiole growth is a key contributor to upward repositioning of the leaf (Polko et al. 2012; Rauf et al. 2013). Intriguingly, shade treatments lead to higher ABA levels in tomato and based on the LRFR-induced gene expression pattern this may also be the case in Arabidopsis (Cagnola et al. 2012; Kohnen et al. 2016). We found that LRFR leads to a rapid increase in ABA content in leaves and therefore decided to study how LRFR controls leaf ABA levels and how ABA controls hyponasty.

## Results

### In LRFR PIFs enhance *NCED* expression and ABA production

Before testing the potential role of ABA in LRFR-induced hyponasty, we first determined whether a reduction of the R/FR ratio (LRFR) rapidly changes ABA levels in Arabidopsis rosettes. 14-day-old-plants grown in long days (LD, 16 hours light, 8 hours darkness), where either kept in standard conditions (white light, WL) or transferred to LRFR (WL supplemented with FR light) at ZT3. Rosettes were collected at the time of transfer to LRFR (ZT3) and 2 hours later (ZT5) and ABA was quantified. This experiment showed that a 2-hour LRFR treatment led to a significant increase in ABA content (Fig. 1A). Since ABA levels fluctuate during the day (Lee et al. 2006; Adams et al. 2018), we also tested the effect of the LRFR treatment later in the day and found that ABA levels increase both in WL and LRFR, but at each time point there was more ABA in LRFR than in WL (Fig. S1A). LRFR led to higher ABA levels both in entire rosettes and in dissected leaves 1 and 2, which we used to analyze the hyponastic response (Fig. S1B).

**Figure 1.**
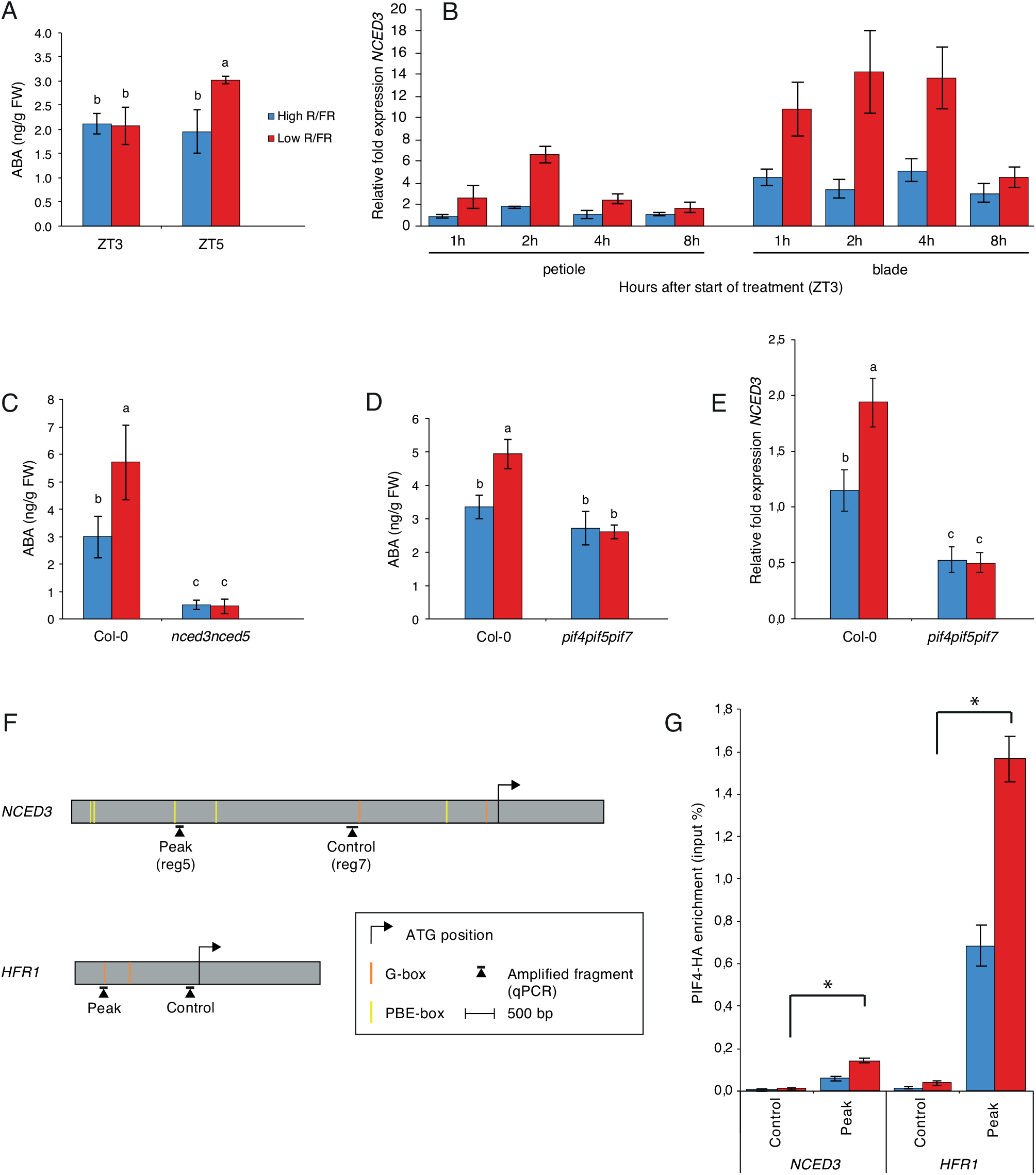
LRFR induces PIF-mediated ABA biosynthesis in leaves. (A) ABA concentration in entire leaves 1 and 2 of wild-type (Col-0) in high R/FR (blue) versus low R/FR (red) conditions at 0 hours (ZT3) and 2 hours (ZT5) after start of treatment. Plants were grown for 14 days in standard long-day [LD, 16-h light, 8-h dark (16/8)] conditions. ZT0 corresponds to the beginning of the light period on day 15. Shade treatment started on day 15 at ZT3 by adding FR light to decrease the R/FR ratio. Each bar plot represents data from 4 biological replicates. Per replicate, 40 leaves (leaves 1 and 2) from 20 plant individuals were harvested and frozen in liquid nitrogen. (B) Relative fold expression of *NCED3* from leaf 3 of Col-0 plants in high R/FR (blue) versus low R/FR (red) conditions over time. Gene expression values were calculated as fold induction relative to a petiole sample at time=1h (ZT4) in high R/FR conditions. Plants were grown for 15 d in standard long-day [LD, 16-h light, 8-h dark (16/8)] conditions. ZT0 corresponds to the beginning of the light period on day 16. Shade treatment started on day 16 at ZT3 by adding FR light to decrease the R/FR ratio. Petioles and lamina of leaf 3 were separately pooled into three biological replicates and frozen in liquid nitrogen. (C) ABA concentration in rosettes of Col-0 versus *nced3nced5* double mutant plants in high R/FR (blue) versus low R/FR (red) conditions at 2 hours (ZT5) after start of treatment. Plants were grown for 14 d in standard long-day [LD, 16-h light, 8-h dark (16/8)] conditions. ZT0 corresponds to the beginning of the light period on day 15. Shade treatment started on day 15 at ZT3 by adding FR light to decrease the R/FR ratio. Each bar plot represents data from 4 biological replicates. Per replicate, 15 and 20 entire rosettes of Col-0 and *nced3nced5* plants, respectively, were harvested and frozen in liquid nitrogen. (D) ABA concentration in rosettes of Col-0 and *pif4pif5pif7* triple plants in high R/FR (blue) versus low R/FR (red) conditions at 2 hours (ZT5) after start of treatment. Plants were grown for 14 d in standard long-day [LD, 16-h light, 8-h dark (16/8)] conditions. ZT0 corresponds to the beginning of the light period on day 15. Shade treatment started on day 15 at ZT3 by adding FR light to decrease the R/FR ratio. Each bar plot represents data from 4 biological replicates. Per replicate, 15 entire rosettes were harvested and frozen in liquid nitrogen. (E) Relative fold expression of *NCED3* in Col-0 and *pif4pif5pif7* from leaves 1 and 2 in high R/FR (blue) versus LRFR (red) conditions. Gene expression values were calculated as fold induction relative to a sample in high R/FR conditions. Plants were grown for 14 d in standard long-day [LD, 16-h light, 8-h dark (16/8)] conditions. ZT0 corresponds to the beginning of the light period on day 15. Shade treatment started on day 15 at ZT3 by adding FR light to decrease the R/FR ratio. At ZT5 (+/− 2h shade), entire leaves 1 and 2 were separately pooled into three biological replicates and frozen in liquid nitrogen. (F) Schematic representation of the *NCED3* and *HFR1* genes. Regions amplified by qPCR and relative positions of G- and PBE-boxes are depicted relative to the start codon. (G) PIF4-HA binding to the *NCED3* and *HFR1* promoter regions. Input and immunoprecipitated DNA was extracted from 10 day-old *PIF4::PIF4-HA(pif4-101*) seedlings exposed to +/− 2 hours of LRFR from ZT2 and quantified by qPCR. PIF4-HA enrichment is presented as IP/input. Bars represent the mean from 3 technical replicates. T-tests were performed and asterisks indicate significant difference (P < 0.05) between peak and control considering shade conditions. (A-E, G) Error bars represent the twofold SE of mean estimates. Two-way ANOVAs followed by Tukey’s Honestly Significant Difference (HSD) test were performed and different letters were assigned to significantly different groups.

The rate-limiting step in ABA biosynthesis is catalyzed by 9-cis-epoxycarotenoid dioxygenase (NCED) enzymes (Qin and Zeevaart 2002). LRFR triggers a rapid increase in *NCED3* and *NCED5* expression in the cotyledons of young seedlings (Kohnen et al. 2016). We therefore tested whether this is also the case in the leaf blades and the petioles of young Arabidopsis rosettes treated with LRFR. *NCED3* and *NCED5* were both rapidly induced in the petiole, while in the leaf blade only *NCED3* expression was induced (Fig. 1B, S1C). Measuring ABA levels in 2-week-old Arabidopsis rosettes showed that ABA levels were strongly reduced in *nced3nced5* double mutants and LRFR did not lead to enhanced ABA accumulation (Fig. 1C). Given the prevalent role of PIF transcription factors for shade-induced transcriptional reprogramming (de Wit et al. 2016), in particular PIF4, PIF5 and PIF7, we compared ABA levels in the wild type (Col-0) and the *pif4pif5pif7* triple mutant. This experiment showed that ABA levels did not increase in *pif4pif5pif7* in response to LRFR (Fig. 1D). To determine whether this is due to PIF-mediated *NCED* expression, we compared the rapid LRFR-induction of *NCED3* in Col-0 and *pif4pif5pif7*. This experiment showed that in *pif4pif5pif7 NCED3* expression was reduced in WL and the rapid LRFR-induction was absent (Fig. 1E).

Genome-wide analysis of PIF4 binding sites identified several PIF4-binding peaks in the *NCED3* promoter of low-blue light grown seedlings (Pedmale et al. 2016). We designed a series of amplicons along the large *NCED3* promoter and found significantly enhanced PIF4 binding in LRFR-grown plants particularly in the region corresponding to peaks 3 and 4 identified in (Pedmale et al. 2016) (Fig. S1D, E). To determine whether the rapid LRFR-induced *NCED3* expression correlates with PIF4 binding, we performed ChIP experiments in plants either maintained in WL or transferred to LRFR for 2 hours and used *HFR1*, a well-known PIF target gene, as a control (Fig. 1F). Our experiments showed enhanced PIF4-HA binding following a LRFR treatment on the promoters of *HFR1* and *NCED3* (Fig. 1G). Collectively these results indicate that PIF transcription factors directly control *NCED3* expression in response to LRFR potentially explaining the observed increase in ABA content upon transfer to LRFR.

### ABA biosynthesis and signaling are required for a normal LRFR-induced hyponastic response

To determine the role of ABA biosynthesis during LRFR-induced hyponasty, we compared the hyponastic response of the wild type with the *nced3nced5* double mutants over 2 days in plants either remaining in WL or being transferred to LRFR at ZT3 of day 1. We focused on leaves 1 and 2, which possess a similar hyponastic response compared to other leaves at the same developmental stage (Dornbusch et al. 2014; Michaud et al. 2017). *nced3nced5* double mutants had more erect leaves than Col-0 throughout the experiment in WL (Fig. 2A). However, this double mutant showed a limited response to a LRFR treatment (Fig. 2A). The *nced3* and *nced5* single mutants had modest phenotypes (Fig. S2A-C). We also tested the *aba2* mutant which is known to have low ABA levels (Gonzalez-Guzman et al. 2002). The phenotype of *aba2* was very similar to that of *nced3nced5* (Fig. 2A, B), consistent with the importance of ABA for normal leaf positioning in WL and a robust hyponastic response to LRFR. We note that despite having more erect leaves *aba2* and *nced3nced5* mutants displayed a normal amplitude of leaf movement in WL, while their ability to respond to LRFR was severely impaired (Fig. 2C).

**Figure 2.**
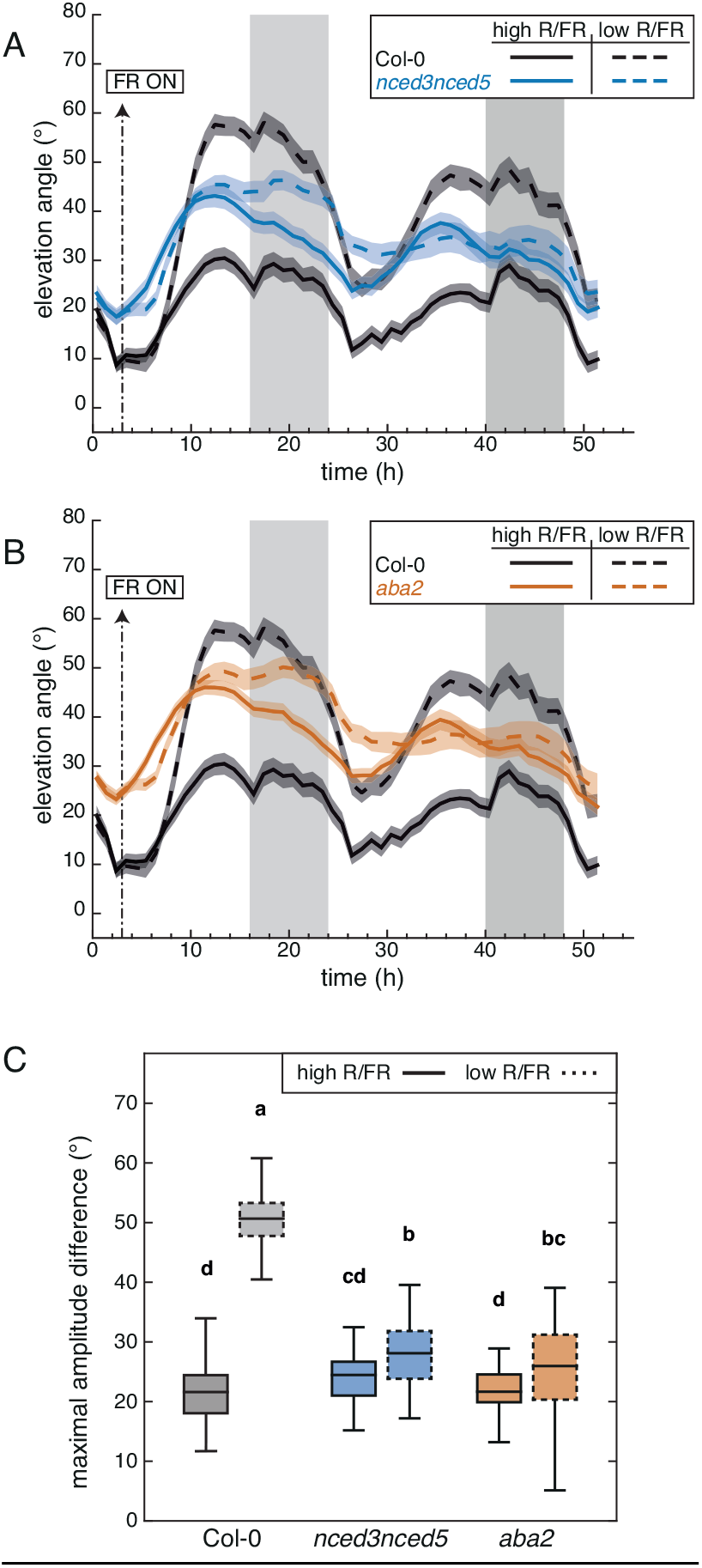
Diel and shade-induced hyponasties both require a functional ABA biosynthetic pathway. (A) Leaf elevation angle of leaves 1 and 2 in Col-0 (black) and *nced3nced5* mutant (blue) plants in high R/FR (solid) versus low R/FR (dashed) conditions. Leaf elevation angles are mean values (n=44-58). (B) Leaf elevation angle of leaves 1 and 2 in Col-0 (black) and *aba2* mutant (orange) plants in high R/FR (solid) versus low R/FR (dashed) conditions. Leaf elevation angles are mean values (n=44-60). (A-B) Plants were grown for 14 days in standard long-day (LD, 16/8) conditions. Imaging started on day 15 at ZT0 (t=0), plants were maintained in LD. Shade treatment started at ZT3 by adding FR light to decrease the R/FR ratio. Col-0 plants analyzed in (A) and (B) are same. Opaque bands around mean lines represent the 95% confidence interval of mean estimates. Vertical gray bars represent night periods. (C) Boxplots representing the amplitude of leaf movement between maximum and minimum leaf elevation angles over the time period from t=3 to t=16 and computed for each individual leaf analyzed in (A) and (B). Solid and dashed plots represent data from high R/FR and low R/FR conditions, respectively. Two-way ANOVA followed by Tukey’s HSD test were performed and different letters were assigned to significantly different groups (p-value < 0.05).

To further investigate the role of ABA in the modulation of leaf hyponasty we analyzed higher order mutants lacking members of the major ABA receptor from the RCAR/PYR1/PYL family (Weiner et al. 2010; Raghavendra et al. 2010). While leaf movements in the quadruple mutant *pyr1pyl1pyl2pyl4* were very similar to Col-0, the *pyr1pyl1pyl2pyl4pyl5pyl8* septuple mutant had an interesting phenotype (Fig. 3A, S3A, B). As observed for ABA biosynthesis mutants, *pyr1pyl1pyl2pyl4pyl5pyl8* had constitutively more erect leaves, a leaf movement amplitude in WL that was at least as robust as in Col-0 but a reduced response to LRFR (Fig. 3A, S3A, B). Higher order mutants lacking protein phosphatases from the ABI1 family of ABA co-receptors (Weiner et al. 2010; Raghavendra et al. 2010) also showed altered leaf movements. This was most striking in the *abi1abi2hab1pp2ca* quadruple (*Qabi2*) mutant, which showed altered leaf movements both in WL and LRFR (Fig. 3B, S3C). In contrast to the other mutants tested, *Qabi2* also showed a strong increase in the leaf elevation angle at night (Fig. S3C). The *hab1abi1pp2ca* triple mutant had wild-type leaf movements in WL but a reduced response to LRFR (Fig. S3D, E). We also analyzed mutants lacking the SnRK2 kinases acting directly downstream of the ABA receptor (Weiner et al. 2010; Raghavendra et al. 2010) and found that the *snrk2.2snrk2.3snrk2.6* triple mutant showed a very severe phenotype (Fig. 3C, S3F). This triple mutant showed constitutively erect leaves with no response to LRFR (Fig. 3C, S3F). This phenotype was only observed in the triple mutant and not in the *snrk2.6* or *snrk2.2snrk2.3* mutants (Fig. S3G). Collectively our data indicate that mutants affecting the positive regulators of ABA signaling (PYR1/PYL and Snrk2 kinases) have very similar phenotypes to ABA biosynthesis mutants. In contrast, mutants affecting members of the ABI1 protein phosphatase family also alter leaf movements and the ability to respond to LRFR but the diel leaf movement patterns were different (Fig. S3).

**Figure 3.**
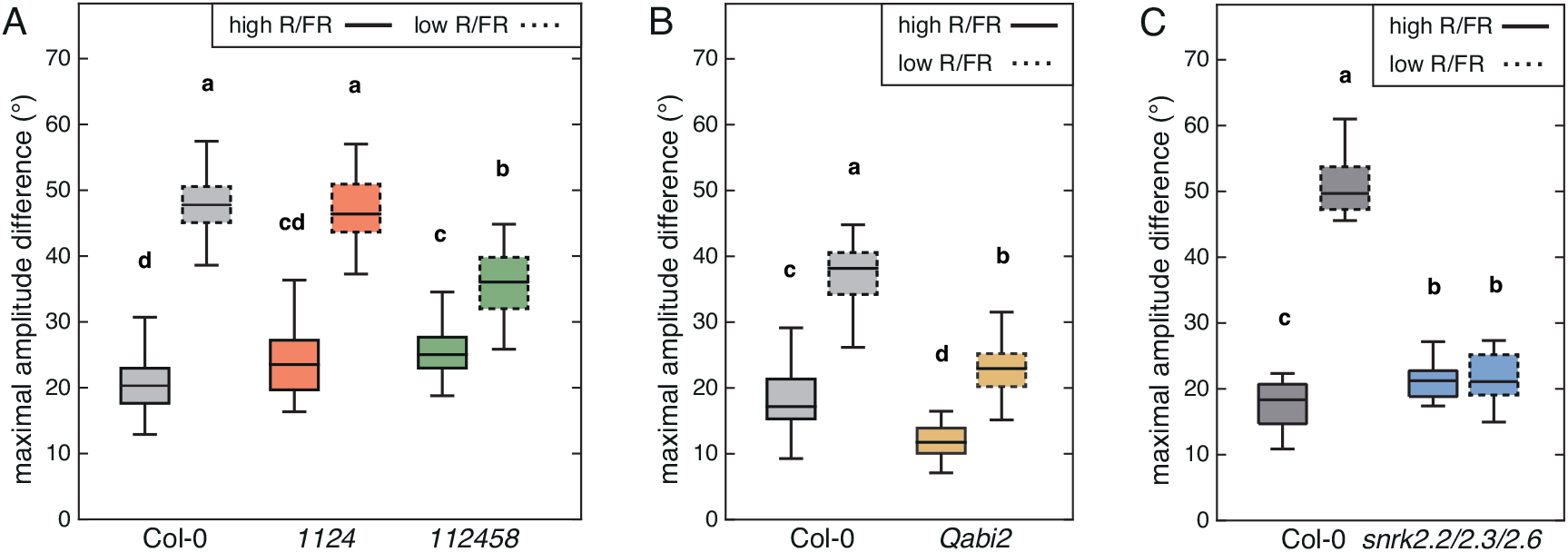
Functional ABA signaling is required for diel and shade-induced hyponasties. (A) Boxplots representing the amplitude of leaf movement between maximum and minimum leaf elevation angles over the time period from t=3 to t=16 of Col-0 (black), *pyr1pyl1pyl2pyl4* quadruple mutant (*1124*, red) and *pyr1pyl1pyl2pyl4pyl5pyl8* sextuple mutant (*112458*, green). Data was computed for each individual leaf analyzed in Figure S4 A, B. Solid and dashed plots represent data from high R/FR and low R/FR conditions, respectively. (B, C) Boxplots representing the amplitude of leaf movement between maximum and minimum leaf elevation angles over the time period from t=3 to t=16 (solid plots, high R/FR) or from t=27 to t=40 (dashed plots, low R/FR) for Col-0 (black), *abi1abi2hab1pp2ca* mutant (B, brown, *Qabi2*) or *snrk2.2snrk2.3snrk2.6* mutant (C, blue) plants and computed for each individual leaf analyzed in Figure S3C (B) or in Figure S3F (C). (A-C) Two-way ANOVA followed by Tukey’s HSD test were performed and different letters were assigned to significantly different groups (p-value < 0.05).

Exogenous auxin (IAA) application to the leaf tip leads to a similar leaf hyponastic response as transferring plants into LRFR (Michaud et al. 2017; Pantazopoulou et al. 2017). We therefore tested whether ABA biosynthesis mutants can respond to auxin application. These experiments showed that *nced3nced5* and *aba2* mutants showed modest responses to IAA application (Fig. 4A, S4A, B). The IAA and LRFR response of these mutants were very similar (Fig. 2–4, S4A, B) suggesting that normal ABA levels are required to respond to the LRFR-induced IAA production required to induce hyponasty. Collectively our data indicate that ABA and ABA signaling modulate diel leaf hyponastic movement and are required for a robust LRFR-induced hyponastic response acting downstream of IAA production.

**Figure 4.**
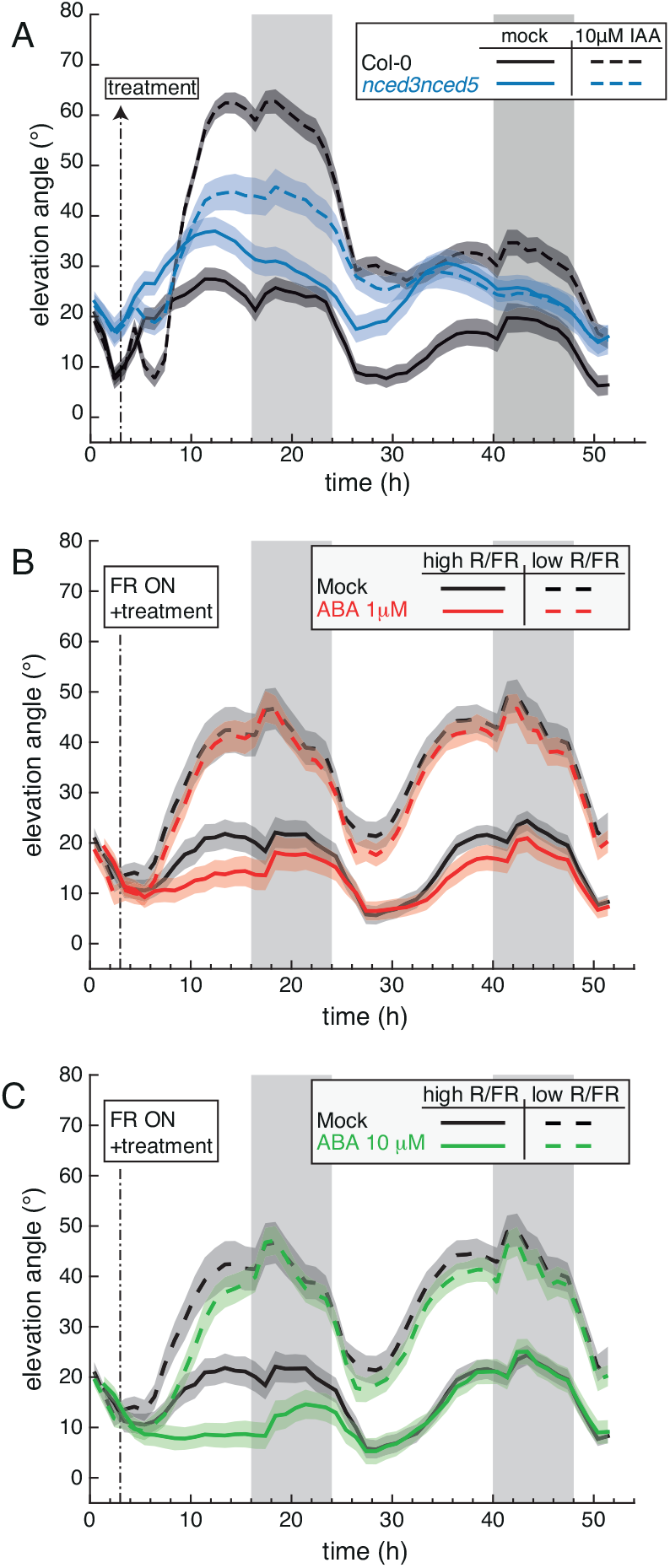
Hormone treatments reveal dysfunctional auxin-induced hyponasty in ABA mutants and reduced sensitivity to ABA in LRFR. (A) Leaf elevation angle of leaves 1 and 2 in Col-0 (black) and *nced3nced5* mutant (blue) plants treated with mock solution (solid lines) or 10μM IAA (dashed lines). At ZT3 on day 15 (t=3) a 1-μL drop of the corresponding solution was applied to the leaf tip (adaxial side). Leaf elevation angles are mean values (n=22-30). (B, C) Leaf elevation angle of leaves 1 and 2 in Col-0 plants treated with mock (black), 1μM ABA (B, red) and 10μM ABA (C, green) solutions in high R/FR (solid) versus low R/FR (dashed) conditions. Leaf elevation angles are mean values (n=26-30). At ZT3 on day 15 (t=3), the corresponding solution was sprayed on the entire leaves (adaxial side). Shade treatment started at t=3 by adding FR light to decrease the R/FR ratio. Col-0 plants analyzed in (B) and (C) are same. (A-C) Plants were grown for 14 days in standard long-day (LD, 16/8) conditions. Imaging started on day 15 at ZT0 (t=0), plants were maintained in LD. Opaque bands around mean lines represent the 95% confidence interval of mean estimates. Vertical gray bars represent night periods.

To determine whether ABA application alters leaf hyponasty we sprayed rosettes with different concentrations of ABA and followed diel leaf movements over 2 days either in WL or LRFR. Consistent with the constitutively high leaf position of WL-grown ABA biosynthesis mutants (Fig. 2), ABA application reduced the leaf angle during the first day after the application in WL in a dose dependent manner (compare Fig. 4B and Fig. 4C). The effect of ABA application was transient as leaf movements of treated and untreated plants were very similar the second day (Fig. 4B, C). Interestingly, in LRFR the effect of exogenous ABA application was largely canceled (Fig. 4B, C). We observed no significant effect following a treatment with 1μM ABA and a delay in the hyponastic response when we applied 10μM ABA (Fig. 4B, C). Our data show that ABA application inhibits leaf elevation happening during diel leaf movements in WL while in LRFR the effect of exogenous ABA was largely canceled suggesting a reduced sensitivity to ABA in shade-mimicking conditions.

### ABA signaling in multiple tissues is required to control hyponasty in LRFR

Given the established importance of ABA in controlling stomata aperture, we first tested whether neighbor-proximity induced changes in leaf transpiration. Young rosettes were transferred from WL to LRFR at ZT3 and a slight increase in transpiration was observed late in the day but the difference between WL and LRFR was not significant (Fig. 5A). We also compared stomata aperture at ZT8 from plants that either remained in WL or were transferred to LRFR at ZT3 and observed no change (Fig. 5B). In contrast, treating plants with ABA or comparing plants during the day with plants that remained in darkness (night extension) validated our stomata opening measuring method as both treatments led to previously documented stomatal closure (Fig. S5A, B). To determine the importance of stomata for diel leaf movements and the response to LRFR we used *epf1epf2* double mutants and EPF2 over-expressing (*EPF2-OX*) plants which maintain a normal stomata morphology but have respectively higher and lower stomata density correlating with higher and lower transpiration (Hepworth et al. 2015). While the *epf1epf2* double mutant had more erect leaves in WL, it responded normally to LRFR (Fig. 5C). *EPF2-OX* plants were very similar to Col-0 (Fig. 5C). Collectively, these data suggest that LRFR-regulated changes in transpiration and stomata aperture are not a major control point for LRFR-induced leaf hyponasty.

**Figure 5.**
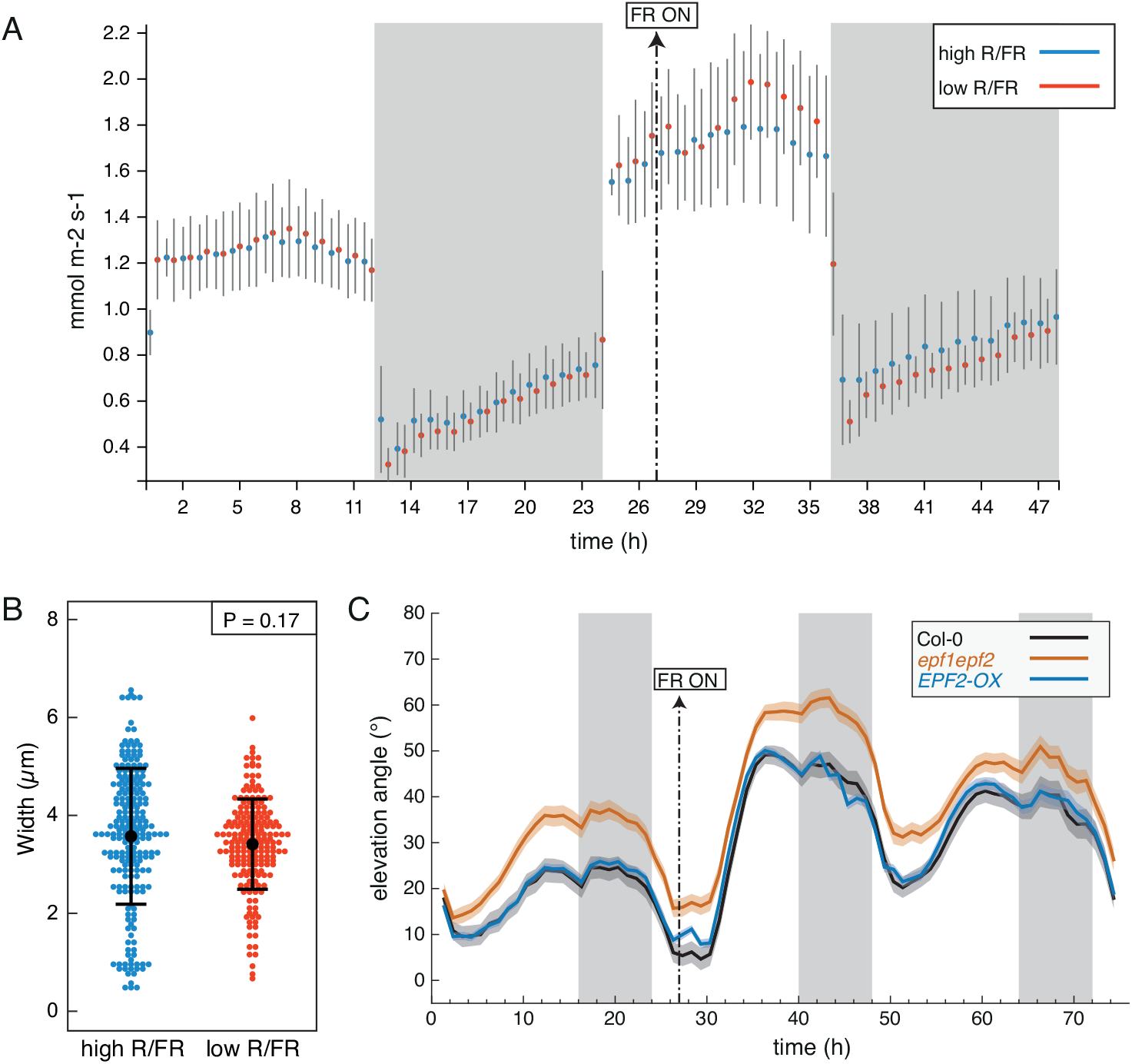
Transpiration and stomata opening during LRFR-induced hyponasty. (A) Transpiration of the aerial parts of 21-day-old Col-0 plants grown in hydroponics system in equinoctial (12:12) conditions, measured by infrared gas analysis (illumination at 150 μmol m^-2^ s^-1^, 20 °C, 60% relative humidity, 380 ppm CO_2_). Mean values from three and four biological replicates (±SD) are given for high R/FR and low R/FR treatments, respectively. Vertical gray bars represent night periods. Supplementation with FR light 3 h into the second day is indicated. (B) Stomatal pore width in high and low R/FR at ZT8 (low R/FR from ZT3), from dental paste imprints of 2-week-old plants grown in LD. P-value given by a T-test without assumption of equal variance. (C) Leaf elevation angle of leaves 1 and 2 in Col-0 (black), *epf1epf2* double mutant (orange), *EPF2-OX* mutant (blue) plants in high R/FR then low R/FR conditions. Shade treatment started on day 16 at t=27 (ZT3) by adding FR light to decrease the R/FR ratio. Leaf elevation angles are mean values (n=29-60). Plants were grown for 14 days in standard long-day (LD, 16/8) conditions. Imaging started on day 15 at ZT0 (t=0), plants were maintained in LD. Opaque bands around mean lines represent the 95% confidence interval of mean estimates. Vertical gray bars represent night periods.

To determine in which cell types ABA signaling is required for normal LRFR-induced leaf hyponasty we generated lines expressing the dominant negative allele *abi1-1* in stomata (*CYP86A2* promoter), mesophyll (*CAB3* promoter) and bundle sheath cells (*SCR* and *MYB76* promoters) (Dickinson et al. 2020). We compared the leaf movements of these plants over 1 long day in WL and then transferred them into LRFR at ZT3 on day 2 and continued imaging for 48 hours. All the lines had a reduced capacity to elevate their leaves in response to LRFR (Fig. 6A-D, S6A-D). The leaf movement pattern of the previously described *pCOR:abi1-1:RFP* line (Duan et al. 2013) was similar to our *pCAB3:abi1-1* lines (Fig. 6A, S6E), consistent with a role of ABA in mesophyll cells to control leaf movements. We also tested the ability of these lines to respond to exogenously applied ABA and found that disrupting ABA signaling in the stomata, mesophyll or bundle sheath cells largely prevented ABA-induced leaf repositioning (Fig. 6E-G, S6F, G). Collectively, our results indicate that different cell types are required for shade-regulated leaf hyponasty and for ABA-induced downward positioning of leaves grown in WL.

**Figure 6.**
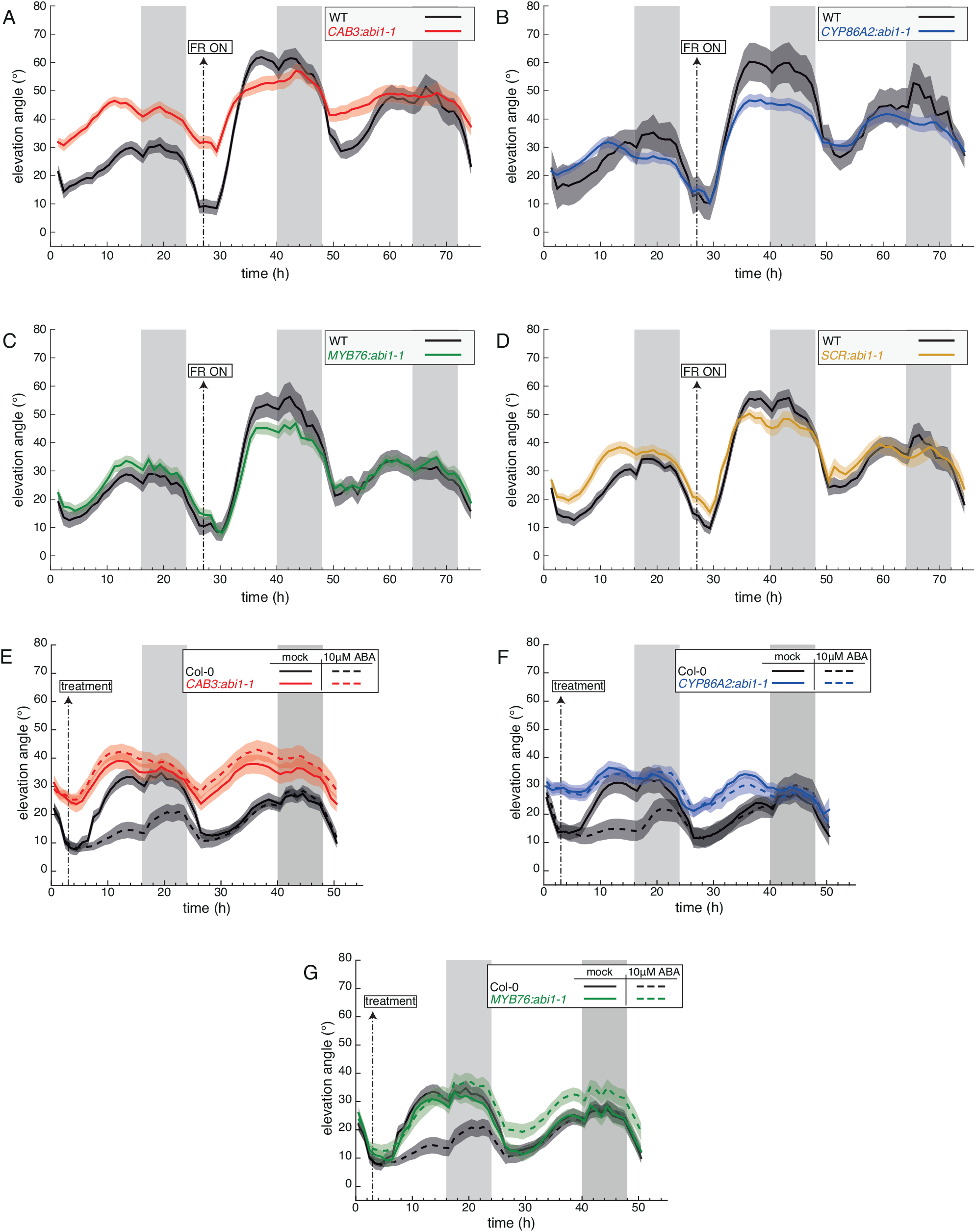
ABA signaling in multiple tissues modulates the hyponastic response. (A-D) Leaf elevation angle of leaves 1 and 2 in Col-0 (A-D, black), *CAB3::abi1-1* mutant (A, red, line #MT36-11), *pCYP86A2::abi1-1* mutant (B, blue, line #MT39-03), *pMYB76::abil-l* mutant (C, green, line #MT41-13), *pSCR::abi1-1* mutant (D, orange, line #MT37-24) plants in high R/FR then low R/FR conditions. Shade treatment started on day 16 at t=27 (ZT3) by adding FR light to decrease the R/FR ratio. Leaf elevation angles are mean values (A, n=18-24; B, n=6-30; C, n=16-28; D, n=16-26). (E-G) Leaf elevation angle of leaves 1 and 2 in Col-0 (E-G, black), *pCAB3::abi1-1* mutant (E, red, line #MT36-11), *pCYP86A2::abi1-1* mutant (F, blue, line #MT39-03), *pMYB76::abi1-1* mutant (G, green, line #MT41-13) plants sprayed with mock solution (solid lines) or 10μM ABA (dashed lines). At ZT3 on day 15 (t=3) mock or ABA solutions were sprayed on the entire rosette (adaxial side). Col-0 plants analyzed in (E) and (G) are same. Leaf elevation angles are mean values (E, n=26-28; F, n=25-28; G, n=21-27). (A-G) Plants were grown for 14 days in standard long-day (LD, 16/8) conditions. Imaging started on day 15 at ZT0 (t=0), plants were maintained in LD. Opaque bands around mean lines represent the 95% confidence interval of mean estimates. Vertical gray bars represent night periods.

## Discussion

We showed that upon transfer to LRFR ABA content rapidly increases in leaves (Fig. 1, S1). Higher ABA levels were also reported in shade treated tomato and in Arabidopsis *phyB* mutants, which display a constitutive shade avoidance response (Cagnola et al. 2012; Gonzalez et al. 2012). Moreover, LRFR leads to increased ABA levels in Arabidopsis buds to prevent lateral shoot development (Gonzalez-Grandio et al. 2013; Holalu and Finlayson 2017). In buds PIFs rapidly induce *BRANCHED1* expression, which was proposed to enhance *NCED3* expression, thereby increasing ABA production (Gonzalez-Grandio et al. 2013; Holalu et al. 2020). In contrast to most previous studies, we looked at a rapid effect of LRFR and detect an increase in *NCED3* and *NCED5* expression within 1-2 hours and higher ABA levels within 2 hours of the shade-simulating treatment (Fig. 1). LRFR promotion of *NCED3* expression and ABA levels depend on PIF4, PIF5 and/or PIF7 and LRFR-induced ABA levels is lost in the *nced3nced5* double mutant (Fig. 1). Consistent with a direct role of PIFs in controlling ABA biosynthesis, we find increased PIF4 binding on the *NCED3* promoter in response to LRFR (Fig. 1, S1). Collectively, our data indicates that in response to LRFR PIFs enhance ABA levels in leaves. In response to LRFR PIFs also mediate rapid IAA production, which, upon asymmetric redistribution in the petiole, controls leaf repositioning (Michaud et al. 2017; Pantazopoulou et al. 2017). Based on IAA application experiments it appears that ABA acts downstream of IAA (Fig. 4, S4). ABA plays a major role in the regulation of water movements (Tardieu et al. 2015; Yoshida et al. 2019). The response to LRFR or temperature elevation triggers rapid IAA production (Quint et al. 2016; Casal and Questa 2018), which promotes growth and repositioning of leaves (van Zanten et al. 2009; Michaud et al. 2017; Pantazopoulou et al. 2017). Whether these responses are due to reversible or irreversible cellular expansion (growth) is not fully understood but in both cases sufficient water supply is required. We propose that in this context, PIF-regulated ABA levels contributes to a normal hyponastic response.

The function of ABA is best understood as a stress hormone. Various abiotic stresses (e.g. drought, salinity) enhance ABA biosynthesis, which limits gas exchange and growth to promote tolerance to unfavorable environments (Yoshida et al. 2019; Kinoshita et al. 2021). For example, when a shade treatment occurs in the presence of high sodium chloride, hypocotyl elongation is inhibited in an ABA-dependent manner (Hayes et al. 2019). However, the relationship between ABA and growth in non-stressful conditions is complex and ABA acts both as an activator or a suppressor of growth (Yoshida et al. 2019). One study on the role of ABA in LRFR-induced hypocotyl elongation concluded that ABA rather prevents growth (Ortiz-Alcaide et al. 2019). The role of ABA in leaf positioning also appears to be complex. In standard light conditions application of ABA prevents hyponasty, while decreasing ABA levels genetically of pharmacologically leads to plants with more erect leaves (Mullen et al. 2006; Benschop et al. 2007) (Fig. 2, 4). In contrast, during high temperature-induced hyponasty ABA was proposed to promote leaf hyponasty (van Zanten et al. 2009). Similarly, we find that in WL conditions mutants with reduced ABA levels or reduced ABA signaling have more erect leaves but maintain robust diel leaf movements (Fig. 2, 3). Applying ABA leads to transient downwards leaf repositioning lasting about a day (Fig. 4). These data indicate that in unstressed conditions ABA limits leaf hyponasty. Having more erect leaves in WL largely correlates with high transpiration. This is true for mutants affecting the PYR/PYL ABA receptors (Gonzalez-Guzman et al. 2012), SnRK2 protein kinases (Fujii and Zhu 2009), ABA biosynthesis mutants (Merlot et al. 2002) or mutants with more stomata (Hepworth et al. 2015). Moreover, PP2C ABA-co-receptor mutants have lower amplitude leaf oscillations during the day in WL (Fig. 3B) and reduced transpiration (Rubio et al. 2009). Collectively this suggests that in WL transpiration, which is controlled by ABA, contributes to leaf position. The role of ABA appears to be different for the LRFR response. Indeed, ABA biosynthesis and the response to ABA are required for full repositioning of the leaves suggesting that in these conditions ABA promotes hyponasty (Fig. 2, 3). Interestingly, the high stomata density mutant *epf1epf2* which, like ABA biosynthesis or ABA response mutants, has more erect leaves in WL and transpires more than the wild type (Hepworth et al. 2015), was able to respond to LRFR (Fig. 5). This suggests that the inability to respond to LRFR in ABA biosynthesis or signaling mutants is not due to their constitutively erect leaves. Moreover, it suggests that having a normal stomata response despite a high or a low transpiration baseline (*epf1epf2* and *EPF2-OX* respectively) (Hepworth et al. 2015), does not prevent LRFR-induced hyponasty (Fig. 5). The different involvement of ABA in controlling hyponasty in high versus low R/FR is also illustrated by the strongly reduced response to applied ABA in LRFR-grown plants (Fig. 4). Similarly, when ABA was applied to heat-treated plants to trigger hyponasty, it did not inhibit upwards repositioning of leaves (van Zanten et al. 2009). Collectively, these experiments show a differential role of ABA in controlling hyponasty during unstressed conditions and in response to changes in light or temperature triggering a strong hyponastic response.

The mechanisms underlying the control of hyponasty by ABA remain poorly understood. In day-night conditions, stomata open rapidly at dawn leading to CO_2_ fixation and enhanced transpiration. However, leaves only start to move upwards several hours later correlating with fast leaf growth (Dornbusch et al. 2014). Hence, stomata opening at dawn does not trigger a rapid leaf elevation in WL conditions. Similarly, when transferred to LRFR, we did not observe significant changes in transpiration or stomatal opening correlating with leaf elevation (Fig. 5, S5). However, based on ABA application experiments (Fig. 4) (Mullen et al. 2006; Benschop et al. 2007) it is reasonable to conclude that regulated stomata opening is required for leaf hyponasty. This is also supported by our finding that *abi1-1* expression from a stomata-specific promoter alters diel leaf movements (Fig. 6, S6). Tissue-specific *abi1-1* expression indicates that ABA signaling is required in multiple tissues for normal leaf positioning (Fig. 6, S6). By controlling transpiration, stomata are a well-known control point of water movements. However, water fluxes are also controlled in other cells including bundle sheath and mesophyll (Negin et al. 2019; Grunwald et al. 2021). In long days, leaf hydraulic conductivity peaks before mid-day (Prado et al. 2019). This pattern in hydraulic conductivity does not directly correlate with leaf position in long-days (see e.g. Fig. 2). In WL, all the *abi1-1* expressing lines were largely unresponsive to ABA application indicating that ABA signaling is required in stomata, mesophyll and bundle sheath to control leaf hyponasty (Fig. S6). Expression of *abi1-1* in mesophyll and stomata clearly altered leaf position in WL, while this was not obvious in the bundle sheath expressing lines (Fig. 6, S6). However, the hyponastic response in plants transferred to LRFR was reduced in all *abi1-1* expressing lines indicating that ABA signaling is required in stomata, mesophyll and bundle sheath for shade-induced leaf hyponasty (Fig. 6, S6). This might be due to altered regulation of the leaf water potential as ABA inhibits H^+^ ATPases in bundle sheath cells similarly to what happens in stomata (Grunwald et al. 2021). It is noteworthy that LRFR also leads to IAA production and a strong auxin response (Michaud et al. 2017; Pantazopoulou et al. 2017). While ABA inhibits H^+^ ATPases, auxin activates these proton pumps (Miao et al. 2022). Hence, we hypothesize that in LRFR the correct balance between IAA and ABA-regulated water movements is required to control leaf positioning. Another striking feature of the LRFR response is the reduced sensitivity to applied ABA (Fig. 4). It is noteworthy that reduced ABA sensitivity in LRFR is predicted based on gene expression in seedlings (Kohnen et al. 2016). Indeed, in cotyledons multiple members of the positively acting *PYR/PYL* and *SNRK2* genes are downregulated, while some of the negatively acting *PP2C* ABA co-receptors genes are upregulated (Kohnen et al. 2016). This could be due to higher ABA levels given that a similar gene expression signature was observed in high light which also triggers ABA production (Huang et al. 2019). That said, high light leads to 4-5 times higher ABA levels typical of a stress response (Huang et al. 2019), while LRFR only led to a 30% increase (Fig. 1). We propose that the mechanisms by which ABA controls leaf position depends on the co-occurrence of an IAA response as observed in LRFR or following an increased temperature. However, the cellular mechanisms by which ABA controls environmentally regulated leaf positioning require future studies.

## Material and methods

### Plant material and plasmid construction

All Arabidopsis lines used in this study are in the Col-0 background, *nced3nced5*, *nced3*, *nced5* (Frey et al. 2012), *aba2* (Leon-Kloosterziel et al. 1996), *pyr1pyl1pyl2pyl4* (Park et al. 2009), *pyr1pyl1pyl2pyl4pyl5pyl8* (Gonzalez-Guzman et al. 2012), *hab1abi1pp2ca* (Rubio et al. 2009), *hab1abi1abi2pp2ca* (named *Qabi2*)(Antoni et al. 2013), *snrk2.2snrk2.3*, *snrk2.6*, *snrk2.2snrk2.3snrk2.6* (Fujii and Zhu 2009), *pif4pif5pif7* (de Wit et al. 2015), *yuc2yuc5yuc8yuc9* (Nozue et al. 2015), *pPIF4::PIF4-3HA (pif4-101*) (Zhang et al. 2017), *epf1epf2*, *EPF2-OX* (Hepworth et al. 2015), *pCOR:abi1-1:RFP* (Duan et al. 2013) were previously described.

All constructs were obtained with the In-Fusion HD cloning kit (Takara) and sequence verified. pCAB3:abi1-1-citrine (pMT36) was obtained by replacing the PHOT1 CDS from *pCAB3:PHOT1-citrine* (Preuten et al. 2013) with *abi1-1* which was amplified with oligonucleotides MT49 and MT50 using plasmid *pEN-1-abi1-1-2* as a template (Barberon et al. 2016). *pSCR:abi1-1citrine* (pMT37) was obtained by replacing the *PHOT1* CDS from *pSCR:PHOT1-citrine* (Preuten et al. 2013) with *abi1-1* which was amplified with oligonucleotides MT49 and MT51 using plasmid *pEN-1-abi1-1-2* as a template (Barberon et al. 2016). *pCYP86A2:abi1-1citrine* (pMT39) was obtained by replacing *pSCR* promoter from pMT37 with the *CYP86A2* promoter which was amplified with oligonucleotides MT57 and MT58 using plasmid *proCYP86A2:GFP-PYR1MANDI* as a template (Park et al. 2015). *pMYB76:abi1-1citrine* (pMT41, 2xDH-35S promoter) was obtained by replacing *pSCR* promoter from pMT37 with the *2xDH-35S* promoter which was amplified with oligonucleotides MT61 and MT62 using plasmid *pMYB76-2xDH-35Smin:GUS* as a template (Dickinson et al. 2020). Transgenic plants were generated by introduction of the plant expression constructs in an *Agrobacterium tumefaciens* strain GV3101/2. Transformation of *A. thaliana* Columbia (Col-0) accession was done by floral dipping. Plasmids contained Basta resistance for plant selection. Oligonucleotides sequences are listed in Table S1.

### Growth conditions and phenotyping analyses

Seeds were stratified at 4°C for 3 d in darkness and then sown on soil saturated with deionized water in a Percival CU-36L4 incubator (Percival Scientific) at 21 °C, 85% relative humidity, and PAR =175 μmol·m^−2^·s^−1^ under LD (16:8) conditions. After 13 days plants were transferred to the ScanAlyzer HTS (LemnaTec) for acclimation (with day–night cycles and light conditions as in the incubator) 24 h before scanning. Experiments were performed under long-day conditions (LD, 16h day/8h night). For shade treatments, the R/FR was decreased from 4.2 to 0.2 using FR-emitting diodes positioned on the ceiling of the ScanAlyzer HTS. Further experimental details, spectral composition of light, computation of the R/FR ratio, and technical specifications of the phenotyping device are described in detail in Dornbusch *et al*. (2012). For gene expression experiments in Fig. 1B and S1C, Col-0 plants were grown as described in de Wit *et al*. (2015). In brief, seeds were directly sown on soil and stratified at 4°C for 3 d in darkness. Plants were then grown for 14 days at 20 °C, 70% relative humidity, and PAR =220 μmol·m^-1^·^−1^ under LD (16:8) conditions. Afterwards, plants were divided over two Percival I-66L incubators (Percival Scientific) at PAR =130 μmol·m^−2^·s^−1^ 24 h before the start of the experiment. Experiment was performed the following day (day 16). At ZT3 on day 16, R/FR was decreased in one of the incubators from 1.4 to 0.2 using FR-emitting diodes positioned on the ceiling.

### Pharmacological treatments

Indole-3-acetic acid (IAA, 10 μM; Sigma-Aldrich) and (+/-)-abscisic acid (ABA, 1-10 μM; Sigma-Aldrich) solutions were freshly prepared from concentrated dimethylsulfoxide (DMSO) and ethanol (EtOH) stocks, respectively, before each application. Mock solutions were similarly prepared to contain 0.15% (v/v) Tween-20, 0.1% (v/v) DMSO and 0.1% (v/v) EtOH, respectively.

### Hormone quantification

ABA measurements were performed as previously described in Glauser *et al*. (2014). In brief, fresh frozen samples were ground to a fine powder using mortars and pestles under liquid nitrogen and about 40 mg of powder was weighed in 2.0 mL Eppendorf tubes. To the tubes were added 5-6 glass beads (2mm diameter), 990 μL of extraction solvent (ethylacetate/formic acid, 99.5: 0.5, v/v) and 10 μL of internal standard solution containing D6-ABA at 100 ng/ml. The tubes were shaken for 4 min at 30 Hz in a tissue lyser, centrifuged for 3 min, the supernatant was recovered and the pellet re-extracted with 500 μL of extraction solvent. Both solutions were then combined, evaporated and reconstituted in 100 μL of methanol 70%. The final extracts were analyzed by UHPLC-MS/MS using an Ultimate 3000 RSLC (Thermo Scientific Dionex) coupled to a 4000 QTRAP (AB Sciex). To quantify ABA in plant samples, a calibration curve based on calibration points at 0.2, 2, 10, 50 and 200 ng/mL, all containing D6-ABA at a fixed concentration of 10 ng/mL, and weighted by 1/x was used (x refers to the concentration of the corresponding calibration point).

### Stomata opening

Stomatal aperture was measured in leaves 1 and 2 of 2-week-old rosettes at ZT8, either in high R/FR or after 5h in LRFR, using dental paste imprints. We applied dental paste (Take 1 Advanced Light Body Wash, Kerr^™^, 34149) to the lower leaf epidermis. After solidification, imprints were removed from the leaf and covered with nail polish (60 Seconds Super Shine 740 Clear, Rimmel, France) which was left to dry and peeled off for visualization by light microscopy. For quantification, ellipses were fit into the opening in ImageJ and the length of the minor axis was used as a measure of stomatal aperture.

### Leaf transpiration

Col-0 plants were grown for 21 d in hydroponics system in equinoctial (12:12) conditions at PAR=150μmol photons m^-2^ s^-1^, 20°C and 60% relative humidity. Growth medium contained 1.5mM KNO_3_, 0.75 mM Ca(NO_3_)_2_, 1 mM MgSO_4_, 1 mM KH_2_PO_4_, 0.5 mM K_2_SO_4_, 0.35 mM CaCl_2_, 12.25 μM MnSO4, 61.25 μM H_3_BO_3_, 875 nM ZnSO_4_, 437.5 nM CuSO_4_ 175 nM (NH_4_)_6_Mo_7_O_24_ and 9 mg/L iron-EDTA (adapted from (Orsel et al. 2004)). Plants were mounted into custom-built gas exchange chambers (Kolling et al. 2015). Three plants were measured in a growth cabinet compartment with a high red to far-red light ratio, i.e. normal growth conditions. Four plants were measured in a growth cabinet compartment with far-red light supplementation by FR-emitting diodes (λ= 740nm). Far-red supplementation was started 3 h after onset of light on the second day. Gas exchange of the aerial parts of the plants was measured continuously for 48 h at 380ppm CO_2_ using an infrared gas analyzer (LI-7000 CO_2_/H_2_O Gas Analyzer, LI-COR Biosciences GmbH). After measurement, leaf area size was extracted from photographs of individual plants and used to normalize the transpiration data. Data was processed, analyzed and plotted using custom-built software (George et al. 2018).

### Analysis of leaf position

For time-lapse experiments, plants were scanned at intervals of 60 minutes with the ScanAlyzer HTS (Lemnatec). As output, we obtained time-lapse images in which the distance of measured plant surface points from a reference plane was color-coded. These images were then transformed into 3D point clouds that yield a precise representation of plant surfaces over time as previously described in Dornbusch *et al*. (2012). Leaf elevation angle (tip elevation angle) was delineated by the vector taking as origin the position of the basal end of the petiole organ and as extremity the position of the tip of the blade organ. A detailed description of the geometric definitions of leaf elevation angle (ϕ_tip_) as well as image and data processing are available in Dornbusch et al. (2012, 2014).

### RT-qPCR

RNA extraction, cDNA reverse transcription and quantitative RT-PCR were performed as previously described (de Wit *et al*. 2015). In brief, entire leaves 1 and 2 (Fig. 1E) or petioles/lamina of leaves 3 (Fig. 1B and S1C, samples from de Wit et al., 2015) were separately pooled into three biological replicates and frozen in liquid nitrogen. After consecutive RNA extraction and reverse transcription, RT-qPCR was performed in three technical replicates for each sample using QuantStudio 6 Flex Real-Time PCR sequence detection system (Applied Biosystems) and FastStart Universal SYBR green Master mix (Roche). Data were normalized against two reference genes (YLS8, UBC) using the Biogazelle qbase software. Gene-specific oligonucleotides used for qPCR reactions are listed in Table S1.

### ChIP-qPCR

Seedlings were harvested and cross-linked as described in (Bourbousse et al. 2012). Subsequent steps of chromatin immune-precipitation were performed as described in (Fiorucci et al. 2020) using an anti-HA antibody (Santa Cruz Biotechnology, Inc., Dallas, TX, USA; sc-7392 X). For the long-term LRFR treatment (Supplementary Figure 1) 4-day-old long-day grown *p35S-PIF4-3XHA* (Lorrain et al. 2008) seedlings were either kept in the same conditions for an additional 3 days or transferred for 3 days into LRFR before harvesting. For the short-term LRFR treatment (Fig. 1) 10-day-old *pPIF4p::PIF4-3XHA* (in *pif4-101*) seedlings were grown in long days and either kept in high R/FR or shifted at ZT2 to LRFR for 2h before harvesting in liquid nitrogen. Oligonucleotides used for ChIP-qPCR reactions are listed in S1 Table.

## Acknowledgements

We thank Gaetan Glauser (Neuchâtel Platform of Analytical Chemistry) for determining ABA content. We thank Annie Marion-Poll (Université Paris-Saclay), Hiroaki Fujii (University of Turku), Julian Schroeder (UCSD), Julie Gray (University of Sheffield), Pedro Rodriguez (IBMCP), José Dinneny (Stanford), Sean Cutler (UC Riverside), Diana Santelia (ETH Zurich) and Niko Geldner (University of Lausanne) for seeds or plasmids.

## Accession numbers

The Arabidopsis Genome Initiative numbers for the genes mentioned in this article are as follows: AT2G18790 (PHYB), AT2G43010 (PIF4), AT3G59060 (PIF5), AT5G61270 (PIF7), AT3G14440 (NCED3), AT1G30100 (NCED5), AT1G52340 (ABA2), AT4G17870 (PYR1), AT5G46790 (PYL1), AT2G26040 (PYL2), AT2G38310 (PYL4), AT5G05440 (PYL5), AT5G53160 (PYL8), AT4G26080 (ABI1), AT5G57050 (ABI2), AT1G72770 (HAB1), AT3G11410 (PP2CA), AT3G50500 (SNRK2.2), AT5G66880 (SNRK2.3), AT4G33950 (SNRK2.6), AT2G20875 (EPF1), AT1G34245 (EPF2), AT4G00360 (CYP86A2), AT1G29910 (CAB3), AT3G54220 (SCR), AT5G07700 (MYB76).

## Supporting Information

**Figure S1.** Detailed analysis of LRFR induced and PIF-mediated ABA biosynthesis in leaves

**Figure S2.** Diel and shade-induced hyponasties in *nced* single mutants

**Figure S3.** Functional ABA signaling is required for diel and shade-induced hyponasties

**Figure S4.** Detailed analysis of auxin-induced hyponasty in ABA biosynthetic mutants

**Figure S5.** Validation of stomatal measurement methodology

**Figure S6.** Dysfunctional ABA signaling in internal leaf tissues affects light-modulated hyponastic responses

**Table S1.** Primers used in this study

