## Supplemental Figures and table for "Abscisic Acid modulates neighbor proximity-induced leaf hyponasty in Arabidopsis"

**Figure S1.** Detailed analysis of LRFR induced and PIF-mediated ABA biosynthesis in leaves

**Figure S2.** Diel and shade-induced hyponasties in *nced* single mutants

**Figure S3.** Functional ABA signaling is required for diel and shade-induced hyponasties

**Figure S4.** Detailed analysis of auxin-induced hyponasty in ABA biosynthetic mutants

**Figure S5.** Validation of stomatal measurement methodology

**Figure S6.** Dysfunctional ABA signaling in internal leaf tissues affects light-modulated hyponastic responses

**Table S1.** Primers used in this study

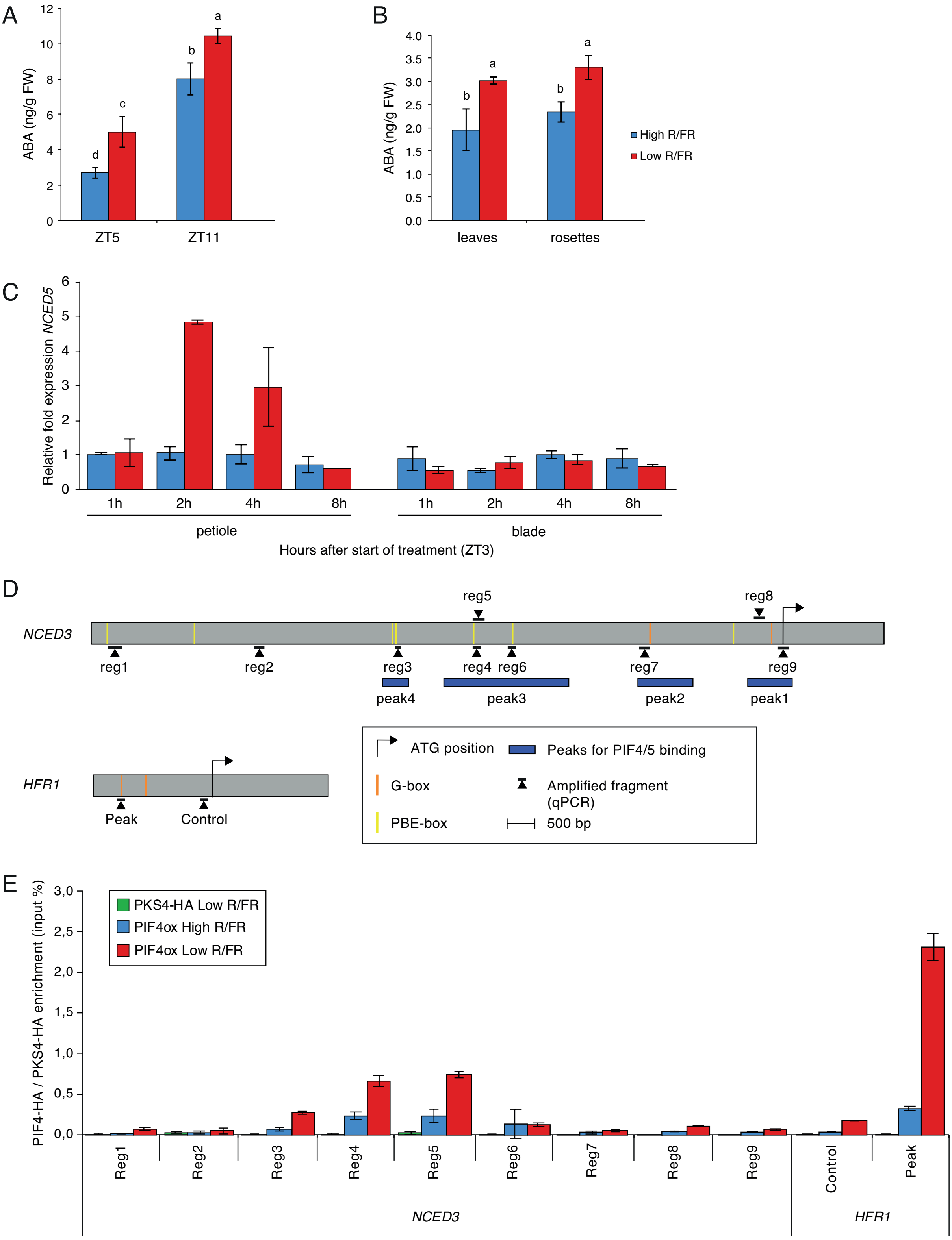


**Figure S1. Detailed analysis of LRFR induced and PIF-mediated ABA biosynthesis in leaves**

(A) ABA concentration in entire leaves 1 and 2 of wild type plants in high R/FR (blue) versus low R/FR (red) conditions at 2 hours (ZT5) and 8 hours (ZT11) after start of the light treatment. Each bar plot represents data from 3 biological replicates. Per replicate, 30 leaves (leaves 1 and 2) from 15 plant individuals were harvested and frozen in liquid nitrogen. (B) ABA concentration in entire leaves (1 and

2) versus entire rosettes of wild type plants in high R/FR (blue) versus low R/FR (red) conditions at 2 hours (ZT5) after start of treatment. Each bar plot represents data from 4 biological replicates. Per replicate, either 40 leaves (leaves 1 and 2) from 20 plant individuals or 10 entire rosettes were harvested and frozen in liquid nitrogen. Data for sampled leaves are the same as presented in Fig.1A. Plants were grown for 14 d in standard long-day [LD, 16-h light, 8-h dark (16/8)] conditions. ZT0 corresponds to the beginning of the light period on day 15. Shade treatment started on day 15 at ZT3 by adding FR light to decrease the R/FR ratio. (A, B) Two-way ANOVAs followed by Tukey's Honestly Significant Difference (HSD) test were performed, and different letters were assigned to significantly different groups. (C) Relative fold expression of *NCED5* from leaf 3 of Col-0 plants in high R/FR (blue) versus low R/FR (red) conditions over time. Gene expression values were calculated as fold induction relative to a petiole sample at time=1h (ZT4) in high R/FR conditions. Plants were grown for 15 d in standard long-day [LD, 16-h light, 8-h dark (16/8)] conditions. ZT0 corresponds to the beginning of the light period on day 16. Shade treatment started on day 16 at ZT3 by adding FR light to decrease the R/FR ratio. Petioles and lamina of leaf 3 were separately pooled into three biological replicates and frozen in liquid nitrogen. (A, B) Two-way ANOVAs followed by Tukey's Honestly Significant Difference (HSD) test were performed, and different letters were assigned to significantly different groups. (D) Schematic representation of the *NCED3* and shade-induced PIF-targeted *HFR1* genes. Regions amplified by qPCR, relative positions of G- and PBE-boxes as well as PIF4/5 ChIP-seq binding sites as reported in Pedmale *et al.* (2016) are depicted relative to the start codon. (E) PIF4-HA and PKS4-HA binding to the *NCED3* and *HFR1* promoter regions. Input and immunoprecipitated DNA was extracted from 7 day-old *P35S::PIF4-3HA (pif4-101)* seedlings exposed to +/- 3 days low R:FR from day 4 as well as PKS4-3HA seedlings exposed to + 3 days low R:FR from day 4 and quantified by qPCR. Seedlings were grown in standard long-day [LD, 16-h light, 8-h dark (16/8)] conditions. PIF4-HA enrichment is presented as IP/input. Bars represent the mean from 3 technical replicates. (A-C,E) Error bars represent the twofold SE of mean estimates.

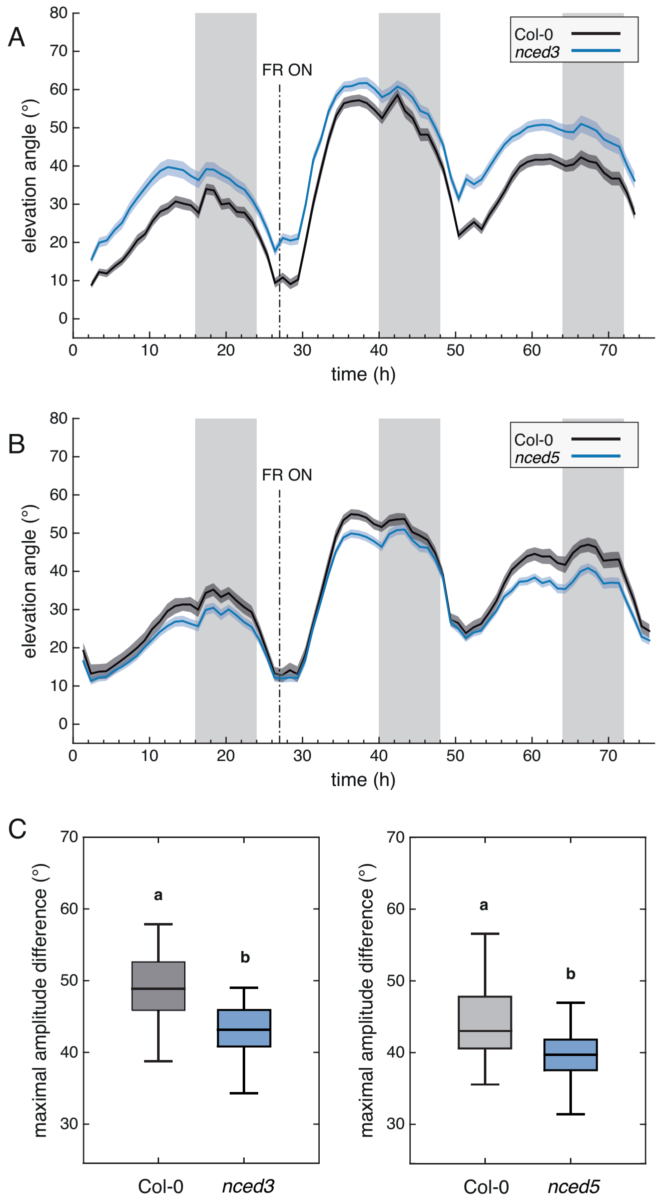


**Figure S2. Diel and shade-induced hyponasties in *nced* single mutants**

(A, B) Leaf elevation angle of leaves 1 and 2 in Col-0 (black) and *nced3* single mutant (A, blue) or *nced5* single mutant (B, blue) mutant plants in high R/FR (t=0-27) then low R/FR conditions (t=27-75). Leaf elevation angles are mean values (A, n=60; B, n= 34-73). Shade treatment started at t=27 (ZT3) by adding FR light to decrease the R/FR ratio. Plants were grown for 14 days in standard long-day (LD, 16/8) conditions. Imaging started on day 15 at ZT0 (t=0), plants were maintained in LD. Opaque bands around mean lines represent the 95% confidence interval of mean estimates. Vertical gray bars represent night periods. (C) Boxplots representing the amplitude of leaf movement between maximum and minimum leaf elevation angles over the time period from t=27 to t=40 and computed for each individual leaf analyzed in (A, left panel) and (B, right panel). Two-way ANOVA followed by Tukey's HSD test were performed and different letters were assigned to significantly different groups (p-value < 0.05).

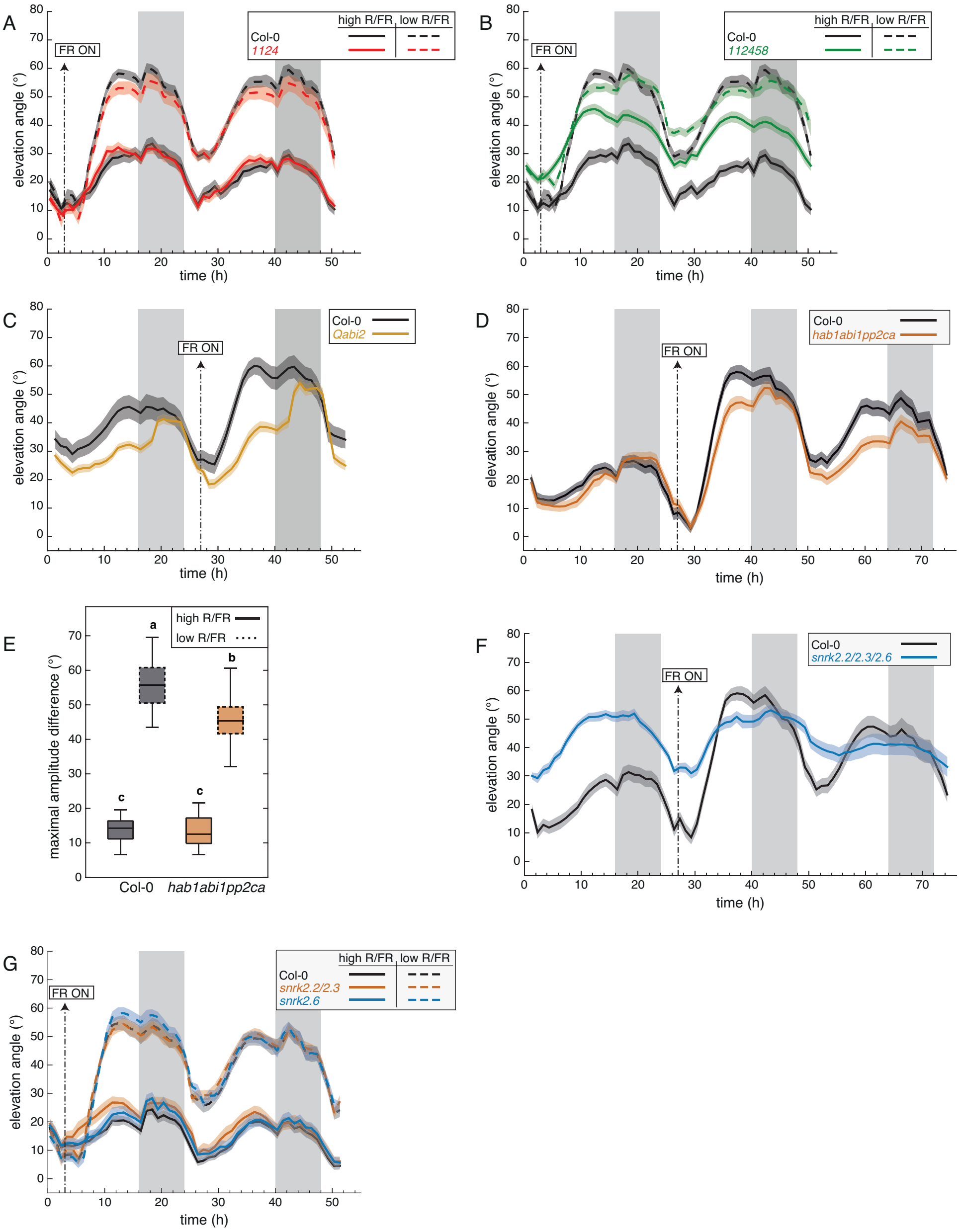


**Figure S3. Functional ABA signaling is required for diel and shade-induced hyponasties**

(A, B) Leaf elevation angle of leaves 1 and 2 in Col-0 (black), *pyr1pyl1pyl2pyl4* quadruple mutant (*1124*, red) and *pyr1pyl1pyl2pyl4pyl5pyl8* sextuple mutant (*112458*, green) plants in high R/FR (solid) versus low R/FR (dashed) conditions. Shade treatment started on day 15 at t=3 (ZT3) by adding FR light to decrease the R/FR ratio. Leaf elevation angles are mean values (n=34-40). Col-0 plants analyzed in (A and B) are the same. (C, D) Leaf elevation angle of leaves 1 and 2 in Col-0 (black), *Qabi2* quadruple mutant (light orange) and *hab1abi1pp2ca* triple mutant (dark orange) mutant plants in high R/FR then low R/FR conditions. Shade treatment started on day 16 at t=27 (ZT3) by adding FR light to decrease the R/FR ratio. Leaf elevation angles are mean values (C, n=23-41; D, n=27-29). (E) Boxplots representing the amplitude of leaf movement between maximum and minimum leaf elevation angles over the time period from t=3 to t=16 (solid plots, high R/FR) or from t=27 to t=40 (dashed plots, low R/FR) and computed for each individual leaf analyzed in (D). Two-way ANOVA followed by Tukey's HSD test were performed and different letters were assigned to significantly different groups (p-value < 0.05). (F) Leaf elevation angle of leaves 1 and 2 in Col-0 (black) and *snrk2.2/2.3/2.6* triple mutant (blue) plants in high R/FR then low R/FR conditions. Shade treatment started on day 16 at t=27 (ZT3) by adding FR light to decrease the R/FR ratio. Leaf elevation angles are mean values (n=22-28). (G) Leaf elevation angle of leaves 1 and 2 in Col-0 (black), *snrk2.2/2.3* double mutant (orange) and *snrk2.6* single mutant (blue) plants in high R/FR (solid) versus low R/FR (dashed) conditions. Shade treatment started on day 15 at t=3 (ZT3) by adding FR light to decrease the R/FR ratio. Leaf elevation angles are mean values (n=26-30). (A-D, F, G) Plants were grown for 14 days in standard long-day (LD, 16/8) conditions. Imaging started on day 15 at ZT0 (t=0), plants were maintained in LD. Opaque bands around mean lines represent the 95% confidence interval of mean estimates. Vertical gray bars represent night periods.

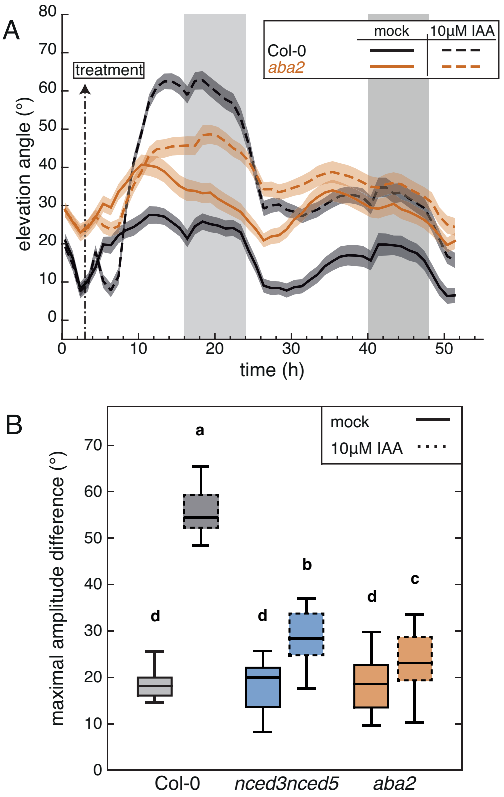


**Figure S4. Detailed analysis of auxin-induced hyponasty in ABA biosynthetic mutants**

(A) Leaf elevation angle of leaves 1 and 2 in Col-0 (black) and *aba2* mutant (orange) plants treated with mock solution (solid lines) or 10µM IAA (dashed lines). At ZT3 on day 15 (t=3) a 1-µL drop of a solution was applied to the leaf tip (adaxial side). Col-0 plants are the same as analyzed in Figure 4A. Leaf elevation angles are mean values (n=24-30). Plants were grown for 14 days in standard long-day (LD, 16/8) conditions. Imaging started on day 15 at ZT0 (t=0), plants were maintained in LD. Opaque bands around mean lines represent the 95% confidence interval of mean estimates. Vertical gray bars represent night periods. (B) Boxplots representing the amplitude of leaf movement between maximum and minimum leaf elevation angles over the time period from t=3 to t=16 and computed for each individual leaf analyzed in Figure 4A (B, *nced3nced5*, blue) and Figure S4A (B, *aba2*, orange). Solid and dashed plots represent data from mock versus IAA treatments, respectively. Two-way ANOVA followed by Tukey's HSD test were performed and different letters were assigned to significantly different groups (p-value < 0.05).

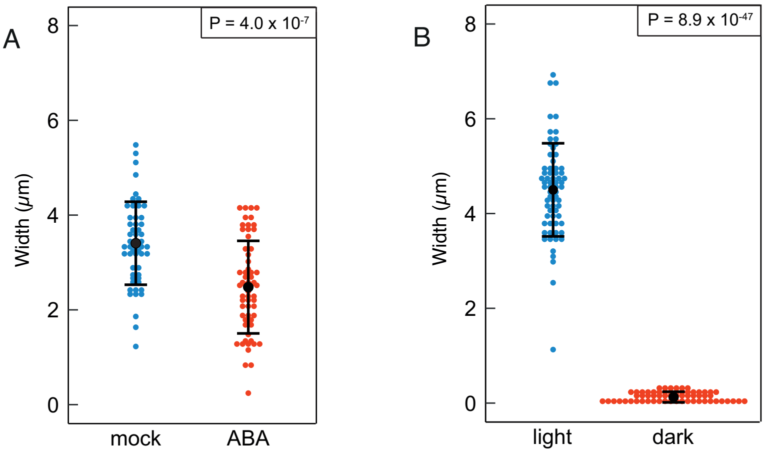


**Figure S5. Validation of stomatal measurement methodology**

(A, B) Stomatal pore widths were measured in conditions that are well known to affect opening, using the same dental paste imprinting approach as in high versus low R/FR in Figure 5. (A) Verification by measuring stomatal opening in light versus darkness: 'light' plants were grown in standard long day conditions and measured several hours after dawn, 'dark' plants were dark adapted from the previous night until measurement of pore width. (B) Verification by measuring stomatal closure in response to ABA treatment 100 µM ABA in 0.1% (v/v) Tween 20. (A, B) P-values given by T-tests without assumption of equal variance.

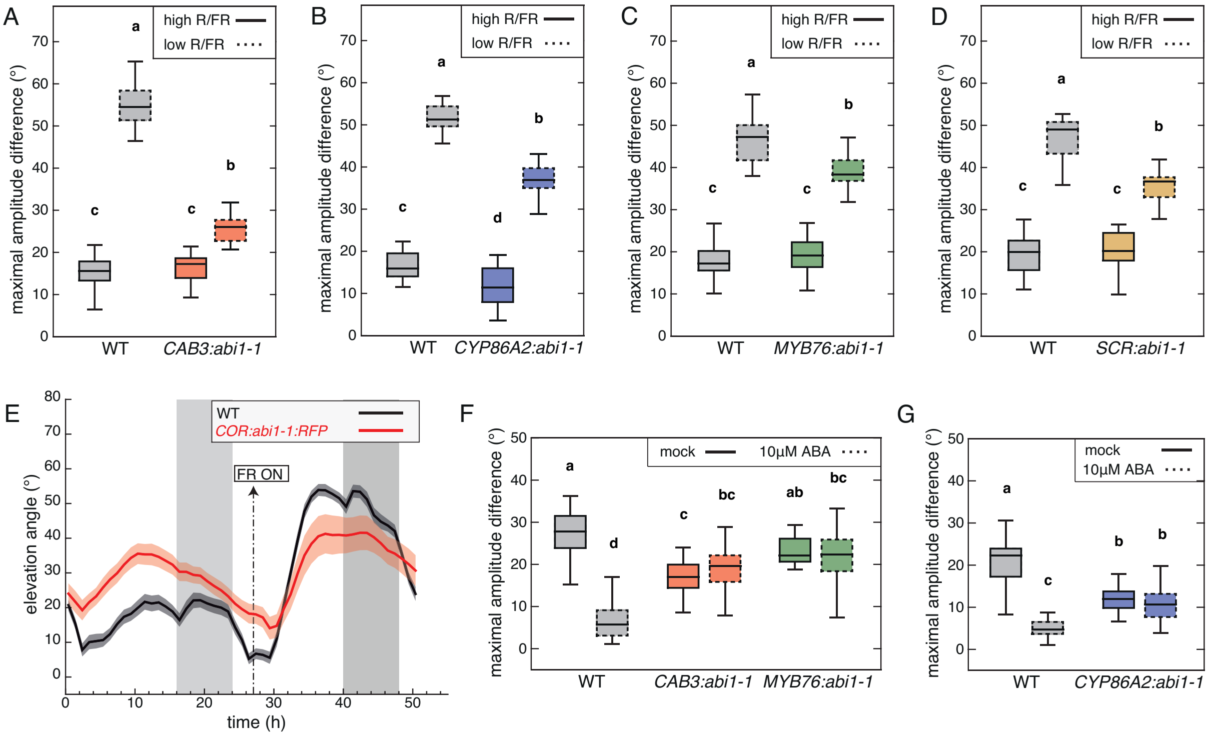


**Figure S6. Dysfunctional ABA signaling in internal leaf tissues affects light-modulated hyponastic responses**

(A-D) Boxplots representing the amplitude of leaf movement between maximum and minimum leaf elevation angles over the time period from t=3 to t=16 (solid plots, high R/FR) or from t=27 to t=40 (dashed plots, low R/FR) and computed for each individual leaf analyzed in Figure 6 A-D, respectively. Two-way ANOVA followed by Tukey's HSD test were performed and different letters were assigned to significantly different groups (p-value < 0.05). (E) Leaf elevation angle of leaves 1 and 2 in Col-0 (black) and *COR::abi1-1::RFP* mutant (red, line #9-2-1) plants in high R/FR then low R/FR conditions. Shade treatment started on day 16 at t=27 (ZT3) by adding FR light to decrease the R/FR ratio. Leaf elevation angles are mean values (n=20). Plants were grown for 14 days in standard long-day (LD, 16/8) conditions. Imaging started on day 15 at ZT0 (t=0), plants were maintained in LD. Opaque bands around mean lines represent the 95% confidence interval of mean estimates. Vertical gray bars represent night periods. (F, G) Boxplots representing the amplitude of leaf movement between maximum and minimum leaf elevation angles over the time period from t=3 to t=16 and computed for each individual leaf analyzed in Figures 6E and 6G (F) as well as in Figure 6F (G). Solid and dashed plots represent data from mock versus 10µM ABA spraying treatments, respectively. Two-way ANOVA followed by Tukey's HSD test were performed and different letters were assigned to significantly different groups (p-value < 0.05).

**Table S1 : oligonucleotides used in this study**

Cloning

MT49: GATATCTTATCTAGAGGATCGTTCAAGGGTTTGCTCTTGAGTTTCC

MT50: AAACACAAAAAGTTTCACCCGGGAAAATGGAGGAAGTATCTCCGG

MT51: CCTTCAATCTCCGGTACCCCGGGAAAATGGAGGAAGTATCTCCGG

MT57: CGGGCCCCCCCTCGAGGACAAGATCACAAGGTTTTCGCATTCAG

MT58: TTCCTCCATTTTCCCGGGTCAATGAATATGAAATGATACTAAAATGGAAAAGTTTAAGGAAT

MT61: CGGGCCCCCCCTCGAGgatgataacacctgaatttaatgacaaaaaaaaaaaaaagtgga

MT62: TTCCTCCATTTTCCCGGGtgttatatctccttggatccgtcgacctgc

Gene expression RT-Q-PCR

NCED3: TCCCTAAGCAATCATCAAACTC / ATTCTTTGGCTTTGGGCTTAAC.

NCED5: GCTCTCATGGCTTGTTCTTAC / GTGAAACTAACGGAGGATGAC.

ChIP-PCR

*NCED3* gene

Reg1: TTCTCTCCGCCAATCCATGA / TTTCTGTCCCACTCTCTCCA (oVCG 591-oVCG 592)

Reg2: GCTCACCACGCAAACACATA / GCAAACTTCTGATGTCGGCA (oVCG 593-oVCG 594)

Reg3: TGTGAATGGTATATCTGAACGCT / TATTGTGCTGTGTGTGGGAC (oVCG 595-oVCG 596)

Reg4: CTCGCGAACCTCCACAAAAT / CGTGCGTAACACATGGAGAC (oVCG 598-oVCG 631)

Reg5: CGTGCGTAACACATGGAGAC / CGGATTTTGGGTCCATAAGA (oVCG 631-oVCG 632)

Reg6: GCACATAGCGTCGGGTTAAA / GACGTGGTTCCATGGTTTCT (oVCG 599-oVCG 600)

Reg7: TCCACCCAAGTTTGCAAATGA / TCCCTCCATCACGATTGCTT (oVCG 601-oVCG 602)

Reg8: TGTCTCGAACCCTCAAACTCA / TGAGACTTGAGACCTTTCACAC (oVCG 605-oVCG 606)

Reg9: ACGATAATGGCGGCTGAGTA / GCCTTTACACATCTCAAAATCG (oVCG 634-oVCG 635)

*HFR1* gene

HFR1 peak: ACGTGATGCCCTCGTGATGGAC / GTCGCTCGCTAAGACACCAAC (PH112-PH113)

HFR1 control: ACGCAACAAACGAACCACAC / AGAGCGATCGGATCAGATAG (PH126-PH127)
